# EuRBPDB: a comprehensive resource for annotation, functional and oncological investigation of eukaryotic RNA binding proteins (RBPs)

**DOI:** 10.1101/713164

**Authors:** Jian-You Liao, Bing Yang, Yu-Chan Zhang, Xiao-Juan Wang, Yushan Ye, Jing-Wen Peng, Zhi-Zhi Yang, Jie-Hua He, Yin Zhang, KaiShun Hu, De-Chen Lin, Dong Yin

**Affiliations:** Guangdong Provincial Key Laboratory of Malignant Tumor Epigenetics and Gene Regulation, Sun Yat-Sen Memorial Hospital, Sun Yat-Sen University, Guangzhou China 510120; Medical Research Center, Sun Yat-Sen Memorial Hospital, Sun Yat-Sen University, Guangzhou, China 510120; Department of stomatology, Sun Yat-Sen Memorial Hospital, Sun Yat-sen University, Guangzhou, China 510120; State Key Laboratory for Biocontrol, School of Life Science, Sun Yat-Sen University, Guangzhou 510275, P. R. China; Department of Medicine, Cedars-Sinai Medical Center, Los Angeles, CA, USA

## Abstract

RNA binding proteins (RBPs) are a large protein family that plays important roles at almost all levels of gene regulation through interacting with RNAs, and contributes to numerous biological processes. However, the complete list of eukaryotic RBPs including human is still unavailable. In this study, we systematically identified RBPs in 162 eukaryotic species based on both computational analysis of RNA binding domains (RBDs) and large-scale RNA binding proteomic (RBPome) data, and established a comprehensive eukaryotic RBP database, EuRBPDB (http://EuRBPDB.syshospital.org). We identified a total of 311,571 RBPs with RBDs and 3,639 non-canonical RBPs without known RBDs. EuRBPDB provides detailed annotations for each RBP, including basic information and functional annotation. Moreover, we systematically investigated RBPs in the context of cancer biology based on published literatures and large-scale omics data. To facilitate the exploration of the clinical relevance of RBPs, we additionally designed a cancer web interface to systematically and interactively display the biological features of RBPs in various types of cancers. EuRBPDB has a user-friendly web interface with browse and search functions, as well as data downloading function. We expect that EuRBPDB will be a widely-used resource and platform for the RNA biology community.

## INTRODUCTION

RNA binding proteins (RBPs) are involved in the regulation of the metabolism, transportation, translation and function of both coding and non-coding RNAs through direct RNA-protein interaction (1). RBPs ensure the smooth flowing of genetic information from DNA to RNA, and ultimately to proteins, making them essential and instrumental for all physiological and pathological processes (1). Numerous diseases have been caused by the aberrant of expression or function of RBPs, including cancer, metabolic disorders and neuropathies (2-4).

Comprehensive identification and annotation of all RBPs are primary and crucial steps for characterization of their functions. To date, several RBPs databases exist for a few eukaryotes, but these databases only collected a small number of well-characterized RBPs from one or few species. For example, RBPDB is a database focusing on the collection of experimentally validated RBPs and RNA binding domains (RBDs), and it contained only 1,171 RBPs from human, mouse, fly and worm (5). ATtRACT is a manually curated database that collects compiled information for only 370 well-characterized RBPs from 39 species (6). Clearly, the RBP repertoire collected by these existing databases are far from complete for any species, human included.

RBPs bind to RNA via structurally well-defined RBDs, such as Dead box helicase domain, RNA recognition motif (RRM) (7,8). Here we annotated proteins containing a RBD as canonical RBPs. Additionally, many studies have suggested the existence of complex protein-RNA interactions that do not require canonical RBDs (9,10), instead through other structures such as intrinsically disordered regions (IDRs) (11). It is thus challenging to identify non-canonical RBPs without known RBDs in a high-throughput and unbiased manner. Recent advances in RNA binding proteome (RBPome) technology significantly facilitate the large-scale identification of non-canonical RBPs (12-18), including the polyT oligos capture RNA interactome method (16-19), click chemistry-based RNA interactome capture (13), and orthogonal organic phase separation (OOPS) of RBPs (14,15,19). These methods crosslink the RBPs with RNA using UV, then apply different strategies to extract total RBPs from cells or tissues. The purified total RBPs are used to analyze the RBPome based on mass spectrometry (MS). These RBPome technologies have been applied to many eukaryotes, including human (15,16,19), mouse (12) and fly (18), and identified a large number of novel canonical and non-canonical RBPs. It should be noted that as a experimental method, none of RBPome technologies is capable of capturing the complete category of RBPs, due to the limitation of total RBP purification strategy and MS technology (12-19).

In the rapid progression of RNA biology field (1), a great need exists to build a comprehensive eukaryotic RBP database to explore the annotation, expression and function of RBPs. To address this, we collected a full list of RBDs from both Pfam (20) and published RBPome datasets from 6 eukaryotes (human, mouse, zebrafish, yeast, fly and worm) (Supplemental Table 1). In parallel, we predicted RBPs based on RBDs using HMMER (21) from the genomes of 162 eukaryotes. Upon integration, we established currently the most comprehensive database of eukaryotic RBP, EuRBPDB (Figure 1). EuRBPDB contains a total of 311,571 RBPs, with detailed annotations for each RBP. Moreover, given the crucial role of RBP in cancer biology, in order to facilitate users to explore the clinical relevance of RBPs, we separately built a Cancer web interface to display integrated cancer-associated omics datasets. The database has a user-friendly interface to interactively exhibit and search the detailed annotations. EuRBPDB will therefore greatly promote the investigation and understanding of the RNA biology.

**Figure 1.**
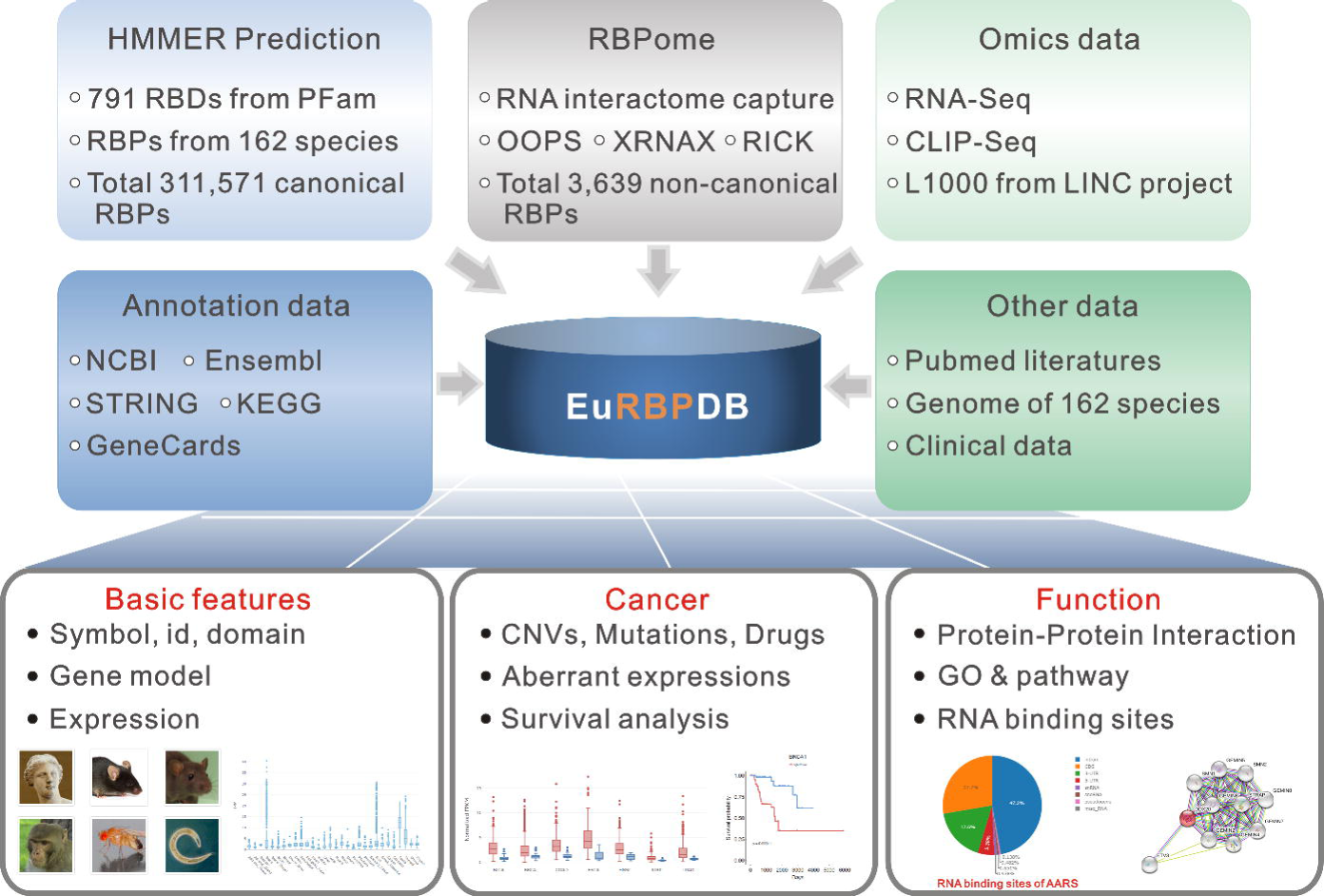
A system-level overview of the EuRBPDB core framework. A total of 315,210 RBPs, including 311,571 canonical RBPs and 3,639 non-canonical RBPs, were identified by combination of computational RBP searching with RBPome profiling. All RBPs were annotated by information retrieved from public database, like NCBI, Ensembl, STRING, KEGG and GeneCards. Cancer-relevant RBPs were identified by literature mining and systematic TCGA data analysis. All the results generated by EuRBPDB were deposited in MySQL relational databases and displayed in the web pages.

## MATERIALS AND METHODS

### Identification and annotation of RBPs

All protein sequences of 162 eukaryotes were downloaded from Ensembl database (release 96, http://www.ensembl.org/). Proteins were annotated as canonical RBPs if they contain one or more domains known to directly interact with RNA. The present RBD list was curated based on the comprehensive RBD list established by Gerstberger et. al.(22). After careful examination, we found that 8 RBDs (RRM6, KH_3, MRL1, Ribosomal_S3_N, Lactamase_B2, tRNA_synt_2b, RnaseH, tRNA_anti) have been removed by Pfam, and thus they were eliminated from our list. Finally, we obtained a total of 791 RBDs (can be downloaded from http://eurbpdb.syshospital.org/data/download/791_RBDs.PFam.gz). We extracted RBD HMM profiles from the Protein families (Pfam) database (Pfam HMM profiles, release v32) (20), and applied the hmmsearch program in HMMER (v3.2.1) (21) package to search for all of the eukaryotic protein sequences against the RBD HMM profiles to identify RBPs. Proteins with E-value less than 0.0001 were considered as bona fide canonical RBPs. In total, we identified 311,571 canonical RBPs from 162 eukaryotic species. In parallel, we manually collected large-scale RBPome datasets of human, mouse, zebrafish, yeast, fly and worm from 20 published works (Supplementary table 1). RBPs presenting in two RBPome datasets were considered to be RBPs of high confidence. As a result, we obtained 3639 non-canonical RBPs from 162 species. Finally, EuRBPDB collected 315,210 RBPs, representing the largest eukaryotic RBP database currently available. The basic information, GO and phenotype annotation of RBPs were obtained from NCBI, Genecards, and Ensembl databases. The protein-protein interaction information was parsed from STRING database (23). The pathway annotation was obtained from KEGG database (24). Expression data were obtained from GTEx (25) and SRA.

### Classification of Eukaryotic RBP family

We characterized and classified canonical RBPs by their sequence-specific RBDs. RBP family was named as the RBD domain if its RBPs only contain one type of RBD. If a RBP contains multiple types of RBDs, it was categorized into the family with the smallest E value in RBD prediction. All non-canonical RBPs were classified as non-canonical RBP family. In total, we obtained 663 RBP families.

### Orthologs and paralogs

The reciprocal best hit (RBH) method (26) was used to predict the putative orthologs of RBPs among different species. We performed the all-against-all BLASTP (v2.7.1+) search between proteins of two genomes with strict cutoffs (E-value ≤ 1e-6, coverage ≥ 50%, identity ≥ 30%) and annotated the reciprocal best hit pairs as orthologs. Paralogs was predicted by the BLAST score ratio (BSR) (27) approach. BLASTP search was conducted in each genome with the same parameters as in orthologs search. The BSR value cutoff was set to 0.4.

### Differential expression, copy number variation (CNV), mutation and survival analysis of RBPs

RNA-Seq, whole-exon sequencing and clinical data were retrieved from TCGA database using R/Bioconductor package TCGAbiolinks (v2.8.4) (28). Differential expression analysis was performed using R package edgeR (v3.22.5) (29) [false discovery rate (FDR) <=1e-5, log2 fold change (log2FC) >=1]. Kaplan-Meier survival analysis was performed by R package survival (v2.43-3). Significant amplification and deletion genomic regions in cancer samples were downloaded from Broad GDAC Firehose website (https://gdac.broadinstitute.org/).

### Cellular effects of drugs to RBP expression

Two L1000 assay level-5 datasets (GSE92742 and GSE70138) (30) generated by the Library of Integrated Cellular Signatures (LINCS) project were downloaded from GEO. These datasets contain over 1,600,000 subdatasets measuring the effects 30,744 drugs on the RNA profiles of 44 cell lines. L1000 assay datasets were parsed and displayed by campR (v1.0.1) and ggplot2 (v3.1.0) R packages as suggested by LINCS project. Expression of RBPs is displayed as z-score.

### RNA binding sites of RBPs

∼270 eCLIP datasets generated from K562 (120 RBPs) and HepG2 (150 RBPs) cell lines were retrieved from ENCODE database (https://www.encodeproject.org/). Peak and bam files of each datasets were downloaded. We used intersectBed of bedtools package (v2.27.1) (31) to annotate each peak, and used coverageBed of bedtools to retrieve the RPM value of each peak.

### Literature analysis of cancer-associated RBP

Literature mining was conducted in geneclip3 (http://ci.smu.edu.cn/genclip3/). In brief, Entrez ids of all RBPs were submitted to geneclip3. Key words of function model of geneclip3 were set as “cancer or tumor”. geneclip3 was run in GeneRIF mode, and will return the PubMed ids of all literatures that study the RBPs in cancers. Then, the information of all literatures was retrieved from PubMed. RBPs reported in 3 cancer-relevant studies were considered to be cancer-associated.

## DATABASE CONTENT AND WEB INTERFACE

### The web-based exploration of RBPs

EuRBPDB provides genome-wide identification of RBPs in large amount of eukaryotic species based on HMMER searching results combined with RBPome datasets analyses. In total, 315,210 RBPs, including 311,571 canonical RBPs and 3,639 non-canonical RBPs, were identified in 162 eukaryotic species. After the systematic annotation of these RBPs, we designed a user-friendly web interface for users to query the database conveniently and interactively. Users can either browse the entire RBP list of any 162 eukaryotes collected in database, or search for any RBP in any eukaryotes of interest. EuRBPDB provides two different ways to browse the data, one is to browse by species, the other is to browse by family defined by RBDs. On the “Species” page, 162 species were classified into 12 categories according to Ensembl taxonomy. To browse the RBP list of each species, users just need to click the species image of interest, and retrieve the detailed RBP information through the following steps: families->family gene list ->single gene annotation. On the “Family” page, EuRBPDB lists all 663 RBP families from 162 eukaryotes. RBP families were ordered by family size in descending order. By clicking the family name, users will get all RBPs grouped by species in this family. Users can also obtain the detailed information of RBP through the following steps: species->gene list ->single gene annotation.

Users can search the specific RBP of interest using the quick search box at the top right corner of navigation bar in any page, the search will return all RBPs in any species matching the searching criteria. To browse the detailed information of any specific RBP, users can specify both the species and RBP name/ID in “Search” page. Both search and browser functions direct users to the detailed information page of any specific RBP. This page comprises of two subpages, namely “Basic Information” subpage and “Cancer Related Information” subpage (only for human RBPs currently). All three subpages consist of a number of information sections constructed by data collected from other published databases. We can readily add any new section to these subpages, and thus it is easy and convenient to update EuRBPDB regularly. In Basic information subpage, EuRBPDB provides basic information including gene structure (Gene Model, Domain section etc.), expression (Expression section), and functional annotation (Pathway, Gene Ontology section etc.). “Cancer Related Information” subpage will be introduced in the following sections.

### Cancer web interface

RBPs contribute extensively and significantly to numerous processes in cancer biology. To facilitate RBP research in cancer, EuRBPDB provides cancer associated annotation of RBPs in Cancer web interface. Through systematic literature mining using geneclip3 (http://ci.smu.edu.cn/genclip3/), we found that a total of 308 RBPs are reported to be associated with human cancers (reported by at least 3 literatures). Among them, 143 RBPs were frequently investigated (reported by more than 20 literatures). Moreover, we conducted differential expression, somatic mutation, CNV, as well as survival analysis based on TCGA data to reveal comprehensively the alterations of RPBs in human cancers. As a result, we identified 1353 RBPs showing aberrant expression in at least one cancer, 2888 RBPs harboring nonsense mutations and missense mutations, 2840 RBPs having frequent genomic deletions or amplifications, and 2885 RBPs exhibiting significant survival correlation. Among these RBPs with cancer-associated alterations, most of them have hitherto not been reported to be associated with any cancers, providing a valuable and novel resource for cancer researchers. EuRBPDB provides the overview of the cancer-associated RBPs in “Cancer” page, as well as the full list of all published and novel cancer-associated RBPs. By clicking the “Details” link of each RBP, users can be redirected to detailed information page of RBP with Cancer Related Information subpage. There are 7 sections in this subpage, showing the literatures investigating selected RBP (Literatures), binding sites of RBP (CLIP), differential expression boxplot (Differential Expression), mutations in RBP (Mutation), copy number variation (CNV), survival analysis (Survival), as well as the expression changes across 44 different cell lines under the treatment of ∼2000 drugs (30).

### RBPredictor web-server for the annotation of eukaryotic RBPs

A web-based tool, RBPredictor, was further developed to assist users to determine whether the protein of interest (from any eukaryote) is a putative RBP. Such RBP prediction is based on the RBD sets used in this study, and we performed hmm-search program in HMMER (v3.2.1) package to determine whether the protein sequence submitted is a putative RBP (22). In “RBPredictor” page, users are only required to input one or multiple protein sequences in fasta format, or submit a fasta file with protein sequences. If an input protein is identified as a putative RBP, RBPredictor will also list all potential RBDs such protein harbors.

## DISCUSSION AND CONCLUSIONS

In this study, we systematically identified eukaryotic RBPs by integrating both large-scale RBPome experimental data and computational RBD identification data. We identified a total of 311,571 high-confident canonical RBPs and 3,639 non-canonical RBPs without known RBDs in 162 eukaryotes. 2,949 RBPs were identified in human with high confidence, including 1,836 canonical RBPs and 1,113 non-canonical RBPs, significantly expanding the human RBP repertoire. Moreover, most human RBPs were found to have cancer-related alterations. We systematically annotated all eukaryotic RBPs in this study, and constructed the most comprehensive eukaryotic RBPs database, EuRBPDB. Through the integration of various large-scale omics data (such as CLIP-Seq, RNA-Seq and L1000 assay), EuRBPDB provides a comprehensive platform to explore the function and cancer-relevance of RBPs. EuRBPDB also provides a framework to systematically identify eukaryotic RBPs based on RBD searching and RBPome data.

Identification of RBP through RBD matching is a highly effective and accurate approach (22). However, recent RBPome studies showed that a large number of proteins without canonical RBDs also bind RNA, and many of them bind RNA through IDRs (11). Therefore, clearly it is insufficient to identify RBPs merely based on RBD searching. On the other hand, RBPome methods are likewise incapable of detecting all RBPs because of the limitation of total RBP purification strategy and relative low sensitivity of MS technology (14,15,19). Thus, presently a comprehensive way to acquire a more complete RBP repertoire is to combine the computational RBP searching with RBPome profiling.

To verify the reliability of RBP dataset we generated, we have cross-checked against all current RBP databases. The results showed that EuRBPDB identified the vast majority of the RBPs (ranging from 90.1% to 100%) across different species collected by other databases (Figure 2), validating the accuracy and consistency of our work. Furthermore, we used the GO annotation to evaluate the robustness and accuracy of our human RBP list. Indeed, we found that 95.3% of canonical RBP and 73.8% of non-canonical RBP were annotated by RNA-related GO terms, such as ‘RNA-binding’, ‘RNA modification’, ‘endoribonuclease activity’. These results together highlight that our RBP identification approach has high accuracy and robust performance. Many databases have been established to aid the research of RNA biology (5,6,32). However, currently no comprehensive RBP database is available for all species. All existing RBP databases focus on the collection and integration of the structure, RBD, RBP binding sites or disease correlation of small amount of well-characterized RBPs in a limited types of eukaryotes, such as RBPDB (5), ATtRACT (6), SpliceAid-F (32), POSTAR2 (33), starBase (34) etc. Compared with these RBP databases, EuRBPDB provides the largest eukaryotic RBP repertoire (315,210 RBPs), the most comprehensive functional and cancer-associated annotation, and an intuitive and easy-to-use web interface. Therefore, EuRBPDB provides a powerful platform to decode the RBP function and regulatory mechanisms.

**Figure 2.**
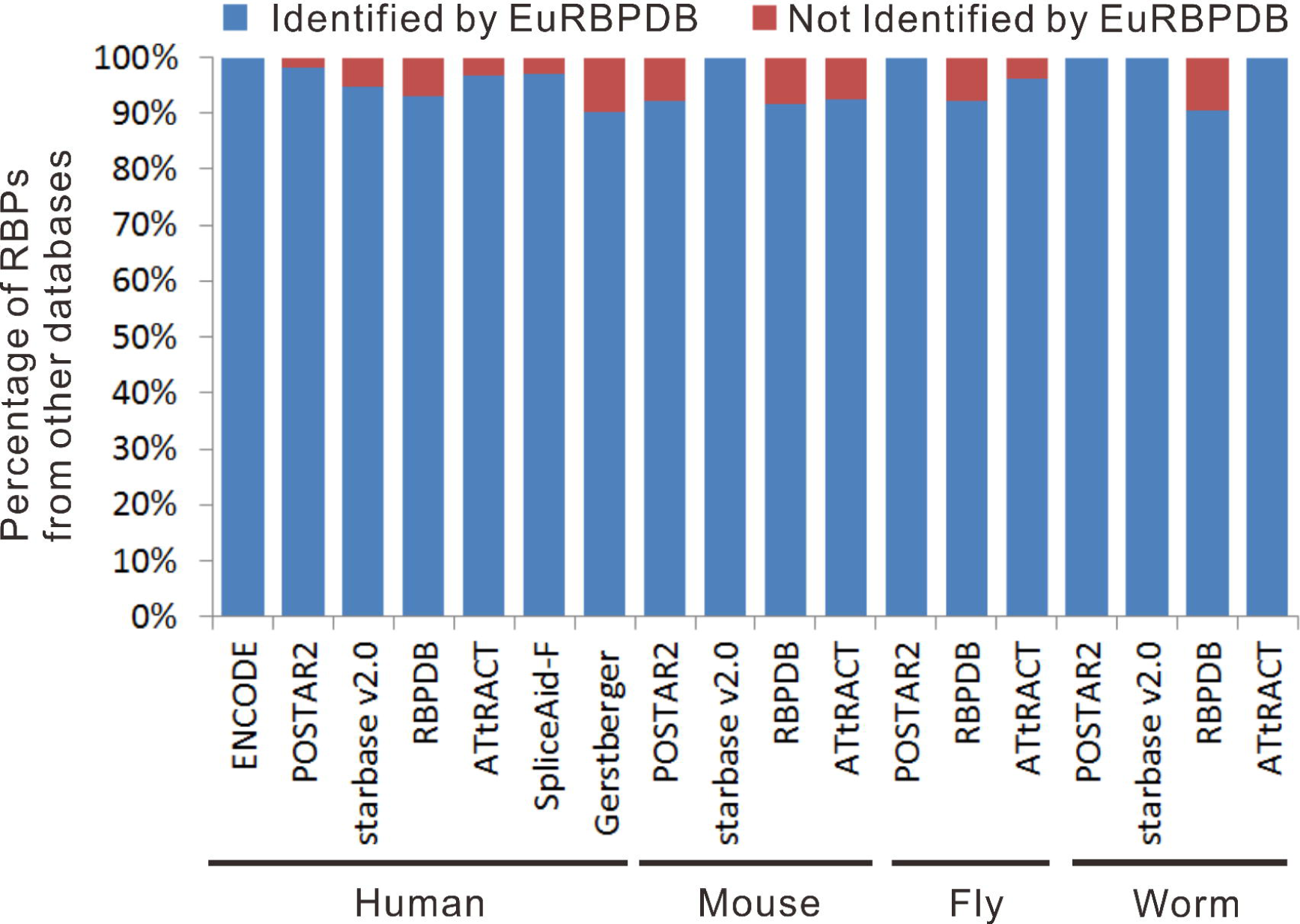
EuRBPDB contained most of RBPs deposited in other databases. 100% (157/157), 98.3% (168/171), 94.7% (36/38), 93.0% (385/414), 96.9% (154/159), 97.0% (65/67), and 90.1% (1,389/1,542) human RBPs from ENCODE database, POSTAR2, starBase v2.0, RBPDB, ATtRACT, SpliceAid-F and Gerstberger et. al. RBP sets (22) respectively were contained in EuRBPDB. 92.31% (36/39), 100% (14/14), 91.7% (373/407) and 92.6% (25/27) mouse RBPs from POSTAR2, starBase v2.0, RBPDB and ATtRACt respectively were included in EuRBPDB. 100% (3/3), 92.2% (226/245) and 96.2% (51/53) Drosophila melanogaster RBPs from POSTAR2, RBPDB and ATtRACT respectively were included in EuRBPDB. 100% (5/5), 100% (2/2), 90.4% (208/230) and 100% (20/20) Caenorhabditis elegans RBPs from POSTAR2, starBase v2.0, RBPDB and ATtRACT respectively were included in EuRBPDB.

## FUTURE DIRECTIONS

EuRBPDB is a comprehensive eukaryotic RBP database, characterizing RBPs of 162 eukaryotic genome-wide. With the ever-increasing amount of RBPome and eukaryotic genome data, we will continue to update and maintain the RBP repertoire and annotation regularly. We will also integrate other types of omics datasets from public databases like Gene Expression Omnibus (GEO) and Sequence Read Archive (SRA) to further improve our understanding of the function and regulatory mechanism of RBPs.

## AVAILABILITY

EuRBPDB database is freely available at http://EuRBPDB.syshospital.org.

